# The proton motive force maintains mtDNA euploidy by balancing mtDNA replication with cell proliferation

**DOI:** 10.1101/2025.10.27.684790

**Authors:** Sneha P. Rath, Vamsi K. Mootha

## Abstract

Human mitochondrial DNA (mtDNA) encodes 13 essential components of the electron transport chain (ETC)^1^. A typical cell contains ∼1000s of copies of mtDNA, but how this copy number is stably maintained is unclear. Here, we track mtDNA copy number (mtCN) recovery in K562 cells following transient, chemically induced depletion to uncover principles of mtCN stability. Below a critical mtCN, ETC activity fails to sustain the proton motive force (PMF) and *de novo* pyrimidine synthesis—both required for mtDNA replication. PMF-dependent processes like Fe-S cluster biogenesis are also disrupted and stress responses are activated that impair cell proliferation and limit further mtCN dilution by cell division. Nonetheless, mtDNA replication and recovery remain possible via mtDNA-independent PMF, generated by complex V reversal, and uridine salvage. Once mtCN is restored, the ETC and forward complex V activity re-engage, stress responses subside, and proliferation recommences. Each cell division then dilutes mtDNA, serving as a built-in brake on mtCN. Our findings suggest that mtCN homeostasis emerges from the balance of two opposing PMF-driven processes — mtDNA replication and cell proliferation — revealing a bioenergetic logic that preserves mtDNA euploidy through repeated cell divisions.

## Introduction

Human mitochondrial DNA (mtDNA) is a small (∼16 Kb), circular, maternally inherited, and high copy number genome^1,2,3^. Proper functioning of mitochondria requires coordination of mtDNA with the nuclear genome: The mtDNA encodes all RNA components of the mitochondrial translation apparatus (2 rRNAs, 22 tRNAs) and 13 proteins which are incorporated into four dual genome-encoded complexes – the proton-pumping electron transport chain (ETC) complexes I, III, and IV, and ATP synthase/complex V (CV). The nuclear genome encodes the remaining 99% of mitochondrial proteins which are synthesized in the cytosol and imported to mitochondria and 15% (200 of 1136)^4^ of these are dedicated to replicating and expressing the mtDNA via mechanisms that have been carefully detailed over the past few decades^5^.

Maintaining the correct number of nuclear and mitochondrial genome copies (euploidy) is critical for cell and organismal fitness. This is widely appreciated for the nuclear genome because aneuploidy is linked to cancer^6^, severe developmental disorders such as Down Syndrome, and miscarriages^7^. Similarly, mtDNA depletion causes clinically heterogenous disorders that impact multiple organ systems and often lead to infant/childhood mortality^8^. MtDNA depletion also impairs proliferation in cell culture, and the knockout of proteins essential for mtDNA replication is embryonic lethal in mice^9^,^10^, further highlighting the essentiality of mtDNA. In contrast, increasing mtDNA copy number (mtCN) buffers against deleterious heteroplasmic mtDNA variants by increasing the total number of wildtype mtDNA molecules^11^.

What has been widely observed is that any given cell type in cell culture will maintain a stable mtCN. Even when mtDNA is chemically depleted, mtCN will return to what appears to be an inherent setpoint for that cell but how this occurs is unclear. In particular, mtDNA maintenance poses a problem of circular dependence because mtDNA replication theoretically depends on products of the mtDNA itself: *de novo* pyrimidine synthesis requires the enzyme DHODH, which is coupled to the mtDNA-encoded ETC^12^, and the proton motive force (PMF), required for import of the mtDNA replication machinery, also depends on the mtDNA-encoded ETC. Yet we know that mtCN can recover even after severe mtDNA loss^13^. How can cells overcome this circular dependence, and what prevents cells from accumulating excess mtDNA, remains to be explained.

The system of mtDNA depletion-recovery serves as an ideal model for understanding mechanisms that ensure mtCN stability. This has typically been achieved by inhibiting mtDNA replication in dividing cells for a few days – using EtBr^14^, inducible mitochondria-targeted nucleases^15^, or a dominant negative POLG^13^ – which depletes mtDNA by dilution with each cell division. When cells are allowed to recover, their mtCN naturally returns to the initial baseline, indicating that cells sense and regulate their intrinsic mtCN. This system allows us to “park” a cell line away from its basal mtCN setpoint, characterize its responses to altered mtCN, and readily examine factors that enable and regulate mtCN (re)establishment.

Here, we combine this depletion-recovery system with multi-omic and bioenergetic analysis and chemical-genetic perturbations to discover that the mtCN is an emergent property that arises from two countervailing processes – mtDNA replication and cell proliferation – that both depend on the proton motive force yet have opposing effects on the mtCN.

## Results

### Proteomic and ultra-structural analysis of a system of reversible mtDNA depletion

To examine the impact of mtDNA loss and how cells sense mtDNA copy number, we leveraged a system of mtDNA depletion-repletion. We treated K562 cells with ethidium bromide (EtBr), which preferentially accumulates in mitochondria, intercalates into mtDNA, and blocks mtDNA replication and transcription. Over five days in EtBr, the cells continued to proliferate without replicating their mtDNA, thus diluting and losing >80% of their mitochondrial genomes. We then grew these mtDNA-depleted cells in EtBr-free media where they resumed mtDNA replication and, after a modest overshoot, re-established their initial mtCN setpoint over 15 days (Figure 1A).

**Figure 1:**
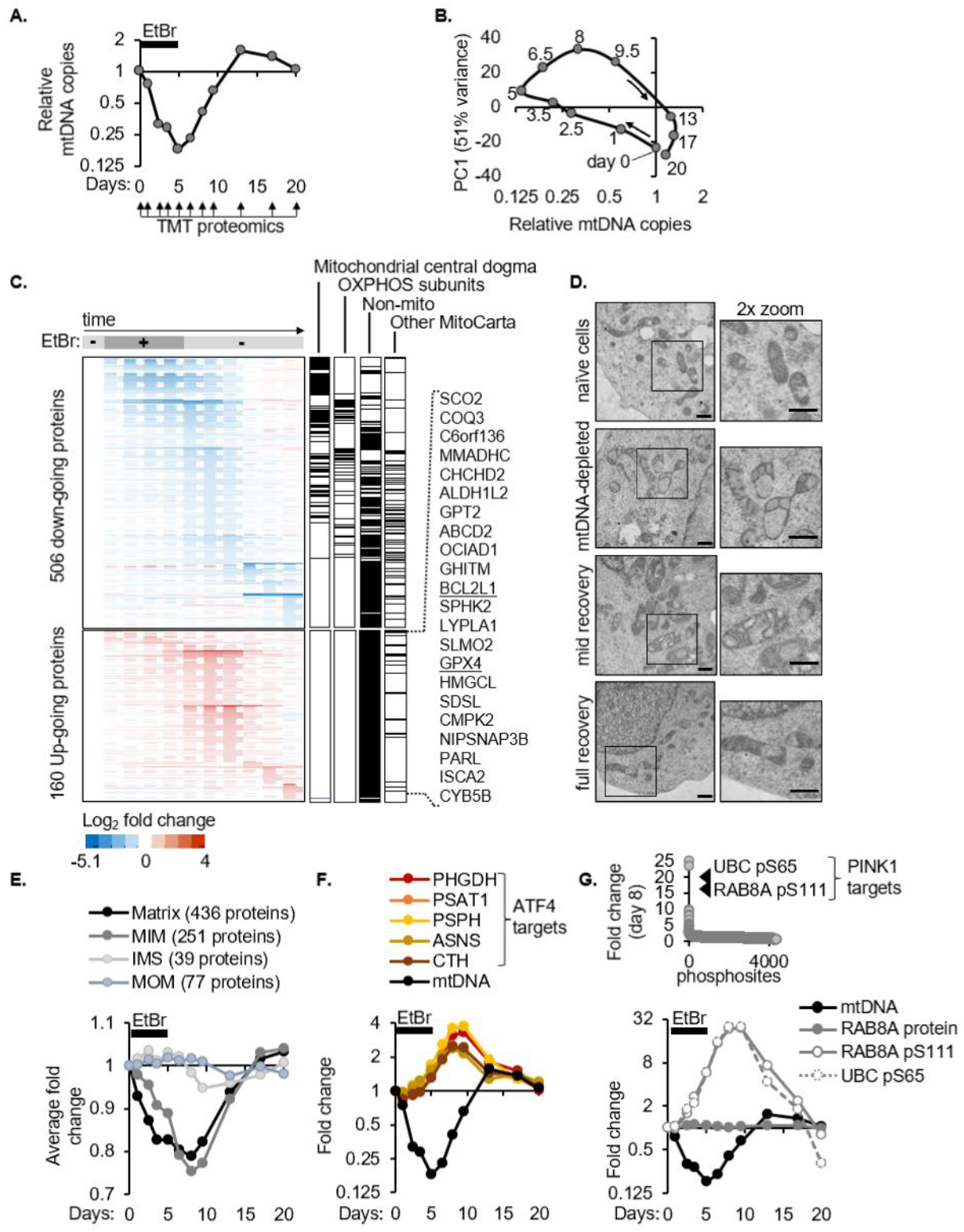
Temporal profiling reveals that mtDNA copy number can recover despite mitochondrial protein loss. **(A)** A system of reversible mtDNA depletion established by treatment with, and subsequent withdrawal of, EtBr. QPCR for mtDNA normalized to nuclear DNA in K562 cells. Eleven samples subject to TMT proteomics from a representative depletion-recovery timecourse. **(B)** Trajectory of PC1 vs mtDNA copy number resulting from PCA analysis of 7,154 proteins detected by TMT proteomics of the mtDNA depletion-recovery timecourse shown in ‘A’. **(C)** Heatmap of all proteins with >1.5-fold change in at least one sample. Dotted boxes show distinct “waves” revealed by hierarchical clustering. Columns of black tick marks indicate genes in each labelled category and where they lie in the heat map. Gene names of the only 22 MitoCarta3.0 proteins that increase during the timecourse are listed on the right. **(D)** Representative images of mitochondria from whole cells imaged by electron microscopy (30,000x). Scale bars show 600 nm with the right column showing a 2x zoom **(E)** Average fold change in all detected mitochondrial proteins categorized by sub-mitochondrial localization. **(F)** Fold change in ATF4 targets in TMT proteomics and qPCR for mtDNA copy number **(G)** Fold change in all detected phosphosites on day 8 of the mtDNA depletion-repletion timecourse (top), full temporal profile of specific phosphosites and total protein, if measured (bottom). Abbreviations: EtBr: ethidium bromide, TMT: Tandem Mass Tag, MIM: mitochondrial inner membrane, IMS: intermembrane space, MOM: mitochondrial outer membrane.

To characterize the consequences of mtDNA loss and define cell states that can support mtDNA replication and euploidy sensing, we performed TMT proteomics at 11 timepoints spanning this 20-day mtDNA depletion-recovery trajectory, complementing our previous transcriptomic characterization of this system^16^. We quantified 7,154 proteins with >1 peptide, including 841 mitochondrial proteins representing 74% of MitoCarta3.0^4^, and several mtDNA-encoded proteins, of which MT-ND3 and MT-CO1 mimic the mtCN overshoot (Figure S1A). Principal component analysis revealed two key features about this system: First, a plot of PC1 vs mtDNA copies showed that majority of the responses to transient mtDNA loss are completely reversible (Figure 1B). Second, there is a “hysteresis” in the system i.e., a lag in the cellular response to changes in mtDNA copy number. For example, day 1 vs day 9.5 have comparable mtDNA copies but are still separated in PC1 (Figure 1B).

The temporal resolution of our TMT proteomics uncovered three major phases of proteome collapse (Figure 1C), with the first phase comprising proteins most proximal to the mtDNA molecule and subsequent phases being increasingly distal. First, concordant with mtDNA loss, is the depletion of exclusively mitochondrial proteins involved in mtDNA maintenance and expression, including the top two proteins positively correlated with steady state mtCN across human cell lines (TFAM and a mitochondrial ribosomal protein)^17^. Second, is the loss of additional players in mtDNA expression as well as oxidative phosphorylation (OXPHOS). Third is the depletion of other mitochondrial as well as non-mitochondrial proteins, and this phase coincides with mtDNA recovery. A “layered” organization had been proposed for the mtDNA nucleoid based on crosslinking experiments^18^ – we now show that the stability of these “layers” is mtDNA-dependent and reversible with mtDNA recovery.

We describe these first two phases of protein collapse through analysis of the five dual-genome encoded complexes – the mitochondrial ribosome and ETC complexes I, III, IV, and V during mtDNA depletion (supplementary figure S1B, C). Across these complexes, two general principles dictate mtDNA-dependent protein stability: i) Subunits physically associated with mtDNA-encoded products deplete to a greater extent with mtDNA loss than distal subunits (Figure S1B, S1C). For example, proteins interacting with mt-rRNAs depleted earlier and more strongly than peripheral proteins which minimally contact mt-rRNAs. (ii) Certain sub-complexes at discrete early steps in the modular assembly of OXPHOS complexes are entirely stable. For example, subunits that nucleate assembly of the Complex I “Q module” and “P_D_-a module” were unchanged in the face of mtDNA depletion(Figure S1C). The collapse of the nuclear-encoded mitochondrial ribosomal and ETC subunits was not observed at the RNA level in previous multi-omic studies of mtDNA depletion^16,19^. Thus, the stability of most nuclear-encoded components within dual-genome encoded complexes is dictated by their mitochondrial RNA/protein partners and assembly intermediates.

While the vast majority of mitochondrial proteins were depleted at some point during the mtDNA depletion-recovery timecourse, interestingly, 22 mitochondrial proteins curiously increase (Figure 1C). We highlight these because they may be under mtDNA-dependent transcriptional/translational control or may be novel candidates like PINK1^20^ or ATFS1^21^ that exhibit mitochondrial import failure and stabilization in a different compartment. We also point out GPX4 and BCL2L1 because their KO is synthetic sick with EtBr^22^, and therefore their upregulation may indicate an adaptive response to cope with mtDNA loss.

### mtCN can recover in cells despite mitochondrial damage and a diminished ETC

Ultrastructure, proteomic, and phospho-proteomic analysis captured the mitochondrial dysfunction ensuing from mtDNA depletion and revealed that it persisted throughout mtDNA recovery. Transmission electron microscopy (TEM) showed that mitochondria of unperturbed K562 cells had parallel, tubular cristae extending deep into the matrix. In contrast, mitochondria of cells with 80% mtDNA loss had large empty pockets in the matrix and barely any cristae (Figure 1D). This aberrant mitochondrial morphology persisted during mtDNA recovery, ultimately reverting to normal only at complete mtCN recovery. This altered morphology and its reversibility was also originally reported in murine L cells by the co-discoverer of mtDNA^23^. Consistent with these TEM images, analysis of mitochondrial proteins by sub-mitochondrial localization^4^ showed that the mitochondrial matrix and inner membrane proteins reached peak depletion on day 8 although EtBr was withdrawn on day 5. (Figure 1E). The ATF4 response, which we previously discovered in the context of mtDNA depletion-recovery^16^, also peaked during mtDNA recovery (Figure 1F). Two *bona fide* PINK1-dependent phosphosites on serine 65 of ubiquitin^24,25^,26 and serine 111 of Rab8A^27^ were the most highly induced out of ∼4000 phosphosites on day 8, indicating mitochondrial depolarization and peak PINK activation. (Figure 1G). Given that mtCN increased between day 5 and day 8, these data collectively indicate that mtDNA replication can be supported in mitochondria with abnormal morphology, bulk protein loss, ISR, and PINK1 activation.

### Residual ETC activity and complex V reversal support recovery of mtCN from its nadir

We would expect mtDNA loss to impair ETC complexes I, III, and IV because their core proton-pumping subunits are mtDNA-encoded. When mtDNA is replete and expressed, the PMF generated by the ETC is used by CV to synthesize ATP (Figure 2A). Upon mtDNA loss, the ETC is diminished and the PMF is generated in an alternative way where CV hydrolyzes ATP in the matrix – also known as CV reversal – to drive an electrogenic exchange of ATP^4-^ with ADP^3-28^ (Figure 2B).

**Figure 2:**
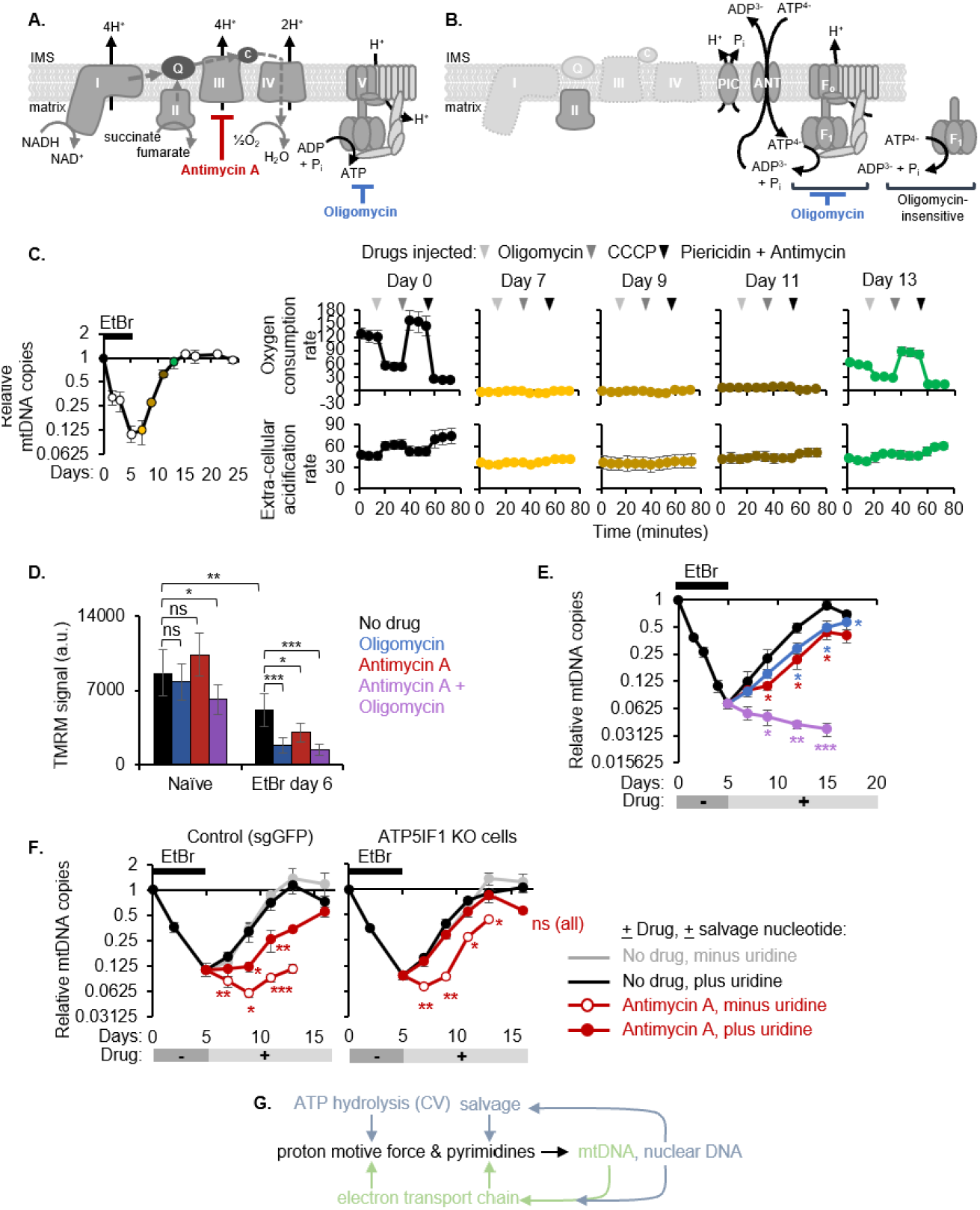
OXPHOS activity is dispensable for mtDNA recovery in cells. **(A)** Schematic of oxidative phosphorylation (OXPHOS) and small molecule inhibitors used in this figure. **(B)** Schematic of OXPHOS upon mtDNA depletion causing loss of complexes I, III, IV and complex V running in reverse (ATP hydrolysis). **(C)** QPCR of mtDNA relative to nuclear DNA normalized to day 0 (left). Filled circles indicate time points for which oxygen consumption and extra-cellular acidification rates from a Seahorse mitochondrial stress test are shown (right, error bars represent standard deviation from five replicates). **(D)** Mitochondrial membrane potential measured by TMRM intensity in mtDNA-depleted (EtBr-treated) or wild type cells after a 30-min exposure to Antimycin A, Oligomycin, or both. Error bars represent standard deviation from six replicates. **(E-F)** QPCR to monitor mtDNA copy number recovery in the presence of the indicated inhibitors and supplements. Error bars represent standard error from three biological replicates. **(G)** Model showing that the proton motive force and pyrimidines required for genome maintenance can be achieved via instructions either from both genomes or solely from the nuclear genome in the case of mtDNA loss or electron transport chain dysfunction. All asterisks indicate p value from a two-tailed Students t-test *<0.05, **<0.01, ***<0.001.

To determine the functional status of the ETC and CV directionality during mtDNA recovery, we used three independent approaches. First, we analyzed the temporal dynamics of each OXPHOS complex by TMT proteomics. The median levels of all OXPHOS complexes continued to decrease past the time point with the lowest mtDNA copy number and recovered with at least a three day-lag with respect to mtDNA (Figure S2A). Nuclear-encoded CV subunits were the most stable of all OXPHOS subunits. This already indicates that the ETC must be dysfunctional during mtDNA recovery due to diminished expression, but CV levels are maintained to allow reversal and defense of the PMF.

Second, we measured oxygen consumption rate (OCR) and extracellular acidification rate (ECAR), which are largely indicative of OXPHOS and glycolysis respectively, in intact cells during mtDNA recovery. While untreated cells had a robust OCR that was sensitive to ETC inhibitors, there was a complete lack of oxygen consumption throughout mtDNA recovery (Figure 2B), consistent with the low protein levels of all ETC complexes. The ETC re-engaged and contributed to oxygen consumption only at complete mtDNA recovery. The ECAR was also insensitive to ETC poisons until complete mtDNA recovery (Figure 2C). Concordantly, the mitochondrial ATP production rate declined with mtDNA depletion and remained negligible during mtDNA recovery. During this period, the glycolytic ATP production rate increased but did not completely compensate for the lost mitochondrial contribution, causing a net deficit in the total ATP production rate (Figure S2B).

Third, we performed an oligomycin setpoint assay to determine how the PMF is defended in the mtDNA-depleted cells that have undetectable OCR. We measured the membrane potential (Δψ), which constitutes the majority of the PMF, using TMRM fluorescence in the presence or absence of acute complex III (CIII) and/or CV inhibition by antimycin and/or oligomycin respectively. Unlike in naïve cells, the PMF of mtDNA-depleted cells collapsed with either antimycin or oligomycin (Figure 2D). The combination of both inhibitors dampened the membrane potential even in naïve cells but did so more strongly in mtDNA-depleted cells. While antimycin directly inhibits CIII, it would also secondarily dampen all other ETC complexes by lowering the oxidized Q pool available for CI/II and by lowering electron flow to CIV.

Thus, our results together indicate that the PMF of mtDNA-depleted cells is maintained by both: residual ETC activity and CV reversal. Having established how the PMF is maintained in the mtDNA-depleted cells in our system, we next probed the bioenergetic requirement for mtDNA replication by tracking mtCN recovery in the presence/absence of antimycin and/or oligomycin. Individually these inhibitors allowed mtDNA recovery with a significant delay and together they prevented mtDNA recovery completely (Figure 2E). This indicates that a certain threshold of the PMF that is jointly defended by residual ETC activity and CV reversal is necessary for supporting mtDNA replication.

### The ETC is dispensable for mtCN recovery via unhindered CV reversal and uridine salvage

MtDNA recovery poses a problem of circular dependence, where mtDNA recovery depends on the ETC, as shown in Fig 2E, but the ETC itself is mtDNA-encoded and can be severely diminished with mtDNA depletion. We aimed to identify a regime where cells can break free from this circular dependence. We hypothesized that in the state of complex III inhibition, when the media already contained glucose and pyruvate to support glycolysis and buffer reductive stress ensuing from ETC dysfunction, the two insults directly preventing mtDNA copy number recovery would be: (i) pyrimidine synthesis halted at the complex III-dependent DHODH reaction and (ii) a drop in the PMF. We sought to rescue each insult by supplementing a salvageable pyrimidine nucleoside, uridine, and knocking out ATP5IF1 to boost the PMF via unhindered CV reversal, respectively. Uridine supplementation alone only partly improves mtDNA recovery of antimycin-treated cells but additionally knocking out ATP5IF1 allows WT-like recovery, regardless of antimycin (Figure 2F).

Thus, pyrimidine synthesis and maintenance of the PMF are two outputs of the ETC that are required for mtDNA recovery but can be alternatively satisfied by nuclear-encoded genes. Thus, the ETC can be rendered completely dispensable for mtDNA maintenance as illustrated in Figure 2G. The PMF can be supported by the mtDNA via the mtDNA-encoded ETC or by the F1 portion of complex V via ATP hydrolysis. Pyrimidine acquisition can also be supported by the mtDNA via the CIII-dependent DHODH reaction or by nucleotide salvage. The latter options for both PMF and pyrimidine acquisition rely exclusively on nuclear-encoded genes and thus alleviate a circular dependence of mtDNA on the protein products it encodes. This redundancy and flexibility in the cell’s ability to defend its PMF and pyrimidine pools is key for maintaining mtDNA in the face of ETC dysfunction.

### Cell proliferation dilutes mtDNA copy number to serve as a brake on mtCN

In dividing cells, we find that proliferation and mtCN regulate each other reciprocally – mtDNA depletion impairs the proliferation of K562 cells^2^ while proliferation effectively dilutes mtCN with each cell division. To decipher the dynamics of this reciprocal regulation, we measured cell proliferation rate throughout our mtDNA depletion-recovery system. We observed that cell proliferation steadily slowed down after the onset of mtDNA depletion, was completely halted during mtDNA recovery, and resumed shortly after mtDNA reached its original copy number set point (Figure 3A, top). The mtCN overshoot and lag in cell proliferation may be due to lingering ATF4 and PINK1 stress responses that resolve only after mtCN recovery (Figures 1F, 1G). We then calculated the rate at which nascent mtDNA molecules had to be synthesized to achieve each steady-state copy number while buffering the dilution effect of cell doubling between time points. In this calculation, we assume no endogenous mtDNA turnover. The mtDNA replication rate was slowest at the onset of recovery from EtBr and improved steadily, reaching its fastest speed when mtDNA copies are replete and overall mitochondrial health returns to the original baseline as shown in figures 1-2 (Figure 3A, bottom). Thus, cells manage to achieve a net recovery of steady-state mtCN, despite a slower initial mtDNA replication rate, because impaired cell proliferation halts the exponential dilution of mtDNA molecules.

**Figure 3:**
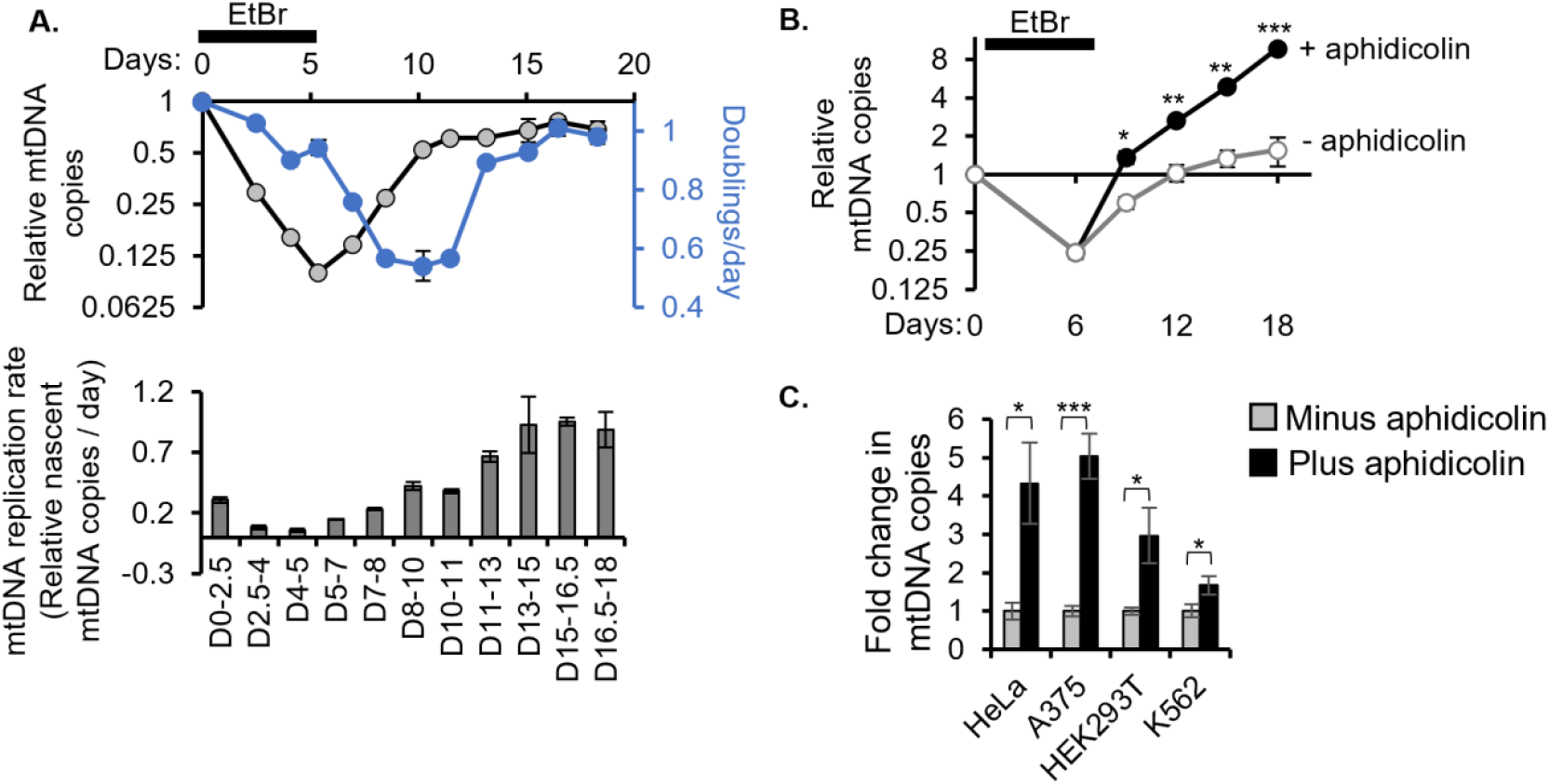
Cell proliferation improves with mtDNA copy number recovery and serves as a built-in brake on mtDNA copy number. **(A)** Steady state mtDNA copy number and cell growth rate (both experimentally measured, top) and mtDNA replication rate (calculated from the first two parameters, see methods, bottom) along the mtDNA depletion-recovery timecourse. **(B)** mtDNA depletion, and recovery in the presence or absence of aphidicolin, quantified by qPCR. **(C)** QPCR for mtDNA copy number in four cell lines treated with or without aphidicolin for 2.5 days. All error bars represent standard error from three replicates and asterisks indicate p value from a two-tailed Students t-test *<0.05, **<0.01, ***<0.001.

We investigated whether cell proliferation slows down during mtDNA recovery due to arrest in a specific cell cycle phase. Cell cycle analysis of asynchronous cells at each timepoint revealed a slight increase in the fraction of cells in G1 during mtDNA recovery (Figure S3A). This partial G1/S arrest may be due to increased phosphorylation of CDK2 at T14/Y15, which is known to be inhibitory^29^ (Figure S3B). G1/S arrest has been reported to occur in a CIV mutant drosophila^30^ and FCCP-treated human cells^31^ via the AMPK/p53 axis. However, given that we do not observe a stark accumulation of cells in any particular phase of the cell cycle despite a near-complete halting of cell proliferation, we infer that mtDNA depletion likely impairs multiple phases of the cell cycle in K562 cells (which are p53 null^32^).

Next, we asked if cell proliferation, which resumes after mtDNA recovery, is itself a brake on the cell’s mtCN setpoint. To prevent cell proliferation from resuming, we treated K562 cells in mtDNA copy number recovery with aphidicolin, which blocks S phase progression. Aphidicolin-treated cells reached an 8-fold higher mtCN than their initial baseline (Figure 3B). Aphidicolin treatment outside the context of mtDNA depletion-recovery also increased the baseline mtCN to varying degrees in three other workhorse cell lines, revealing that cell proliferation limits mtCN (Figure 3C).

### ATP5IF1 KO partially rescues proliferation in mtDNA-depleted cells by rescuing essential mitochondria-dependent processes, such as Fe-S biogenesis

To understand why mtDNA-depleted cells proliferate slowly, we leveraged the only robust genetic suppressor of this EtBr-induced proliferation defect revealed by a previous genome-wide CRISPR KO growth screen^23^ – ATP5IF1, an inhibitor of the ATPase activity of CV’s F1 subunit that is known to boost the growth of mtDNA-depleted cells in multiple organisms^33,34^. In the presence of EtBr or antimycin A and oligomycin (AO), both of which diminish the PMF, ATP5IF1 KO cells exhibited significantly improved proliferation, indicated by increased relative guide abundance over 14 days (Figure 4A). We validated this screening result in polyclonal ATP5IF1 KO K562 cells (Figure S4A). The proliferation defect resulting from chronic EtBr or AO treatment was partially rescued by ATP5IF1 loss (Figure 4B, top). This improved proliferative advantage in the face of EtBr allowed ATP5IF1 KO cells to further deplete their mtCN compared to control cells in EtBr, while AO-treated cells maintained their mtDNA levels regardless of ATP5IF1 status (Figure 4B, bottom).

**Figure 4:**
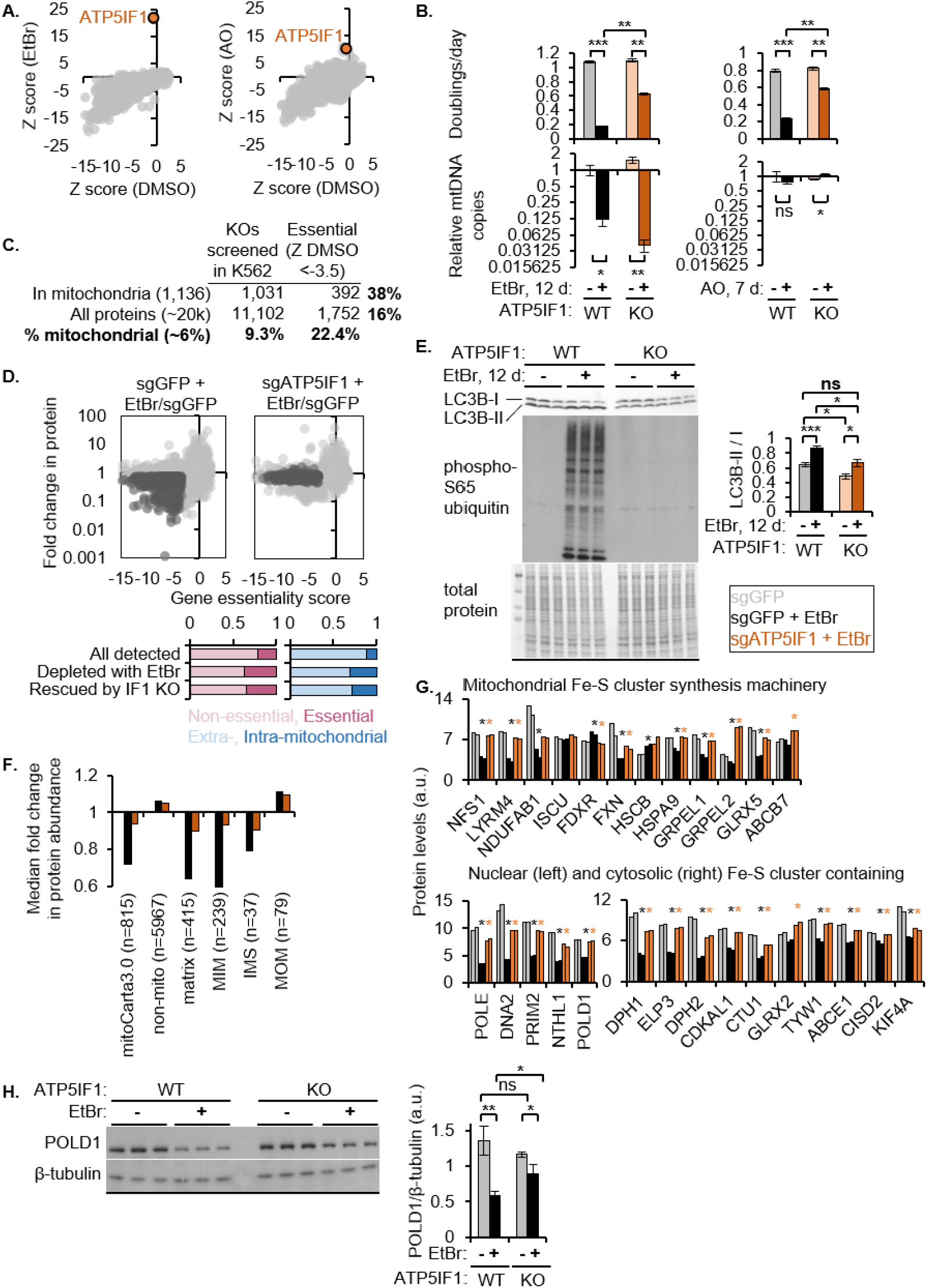
ATP5IF1 KO rescues proliferation and the mitochondrial proteome, including essential processes such as Fe-S cluster biosynthesis, in mtDNA-depleted cells. **(A)** Z score of ATP5IF1 KO in pooled genome-wide CRISPR KO growth screens in the presence of EtBr or AO over 15 days (To *et. al*.) **(B)** Growth rate and mtDNA copy number of control or polyclonal ATP5IF1 KO K562 cells in the presence of EtBr or AO. Growth assessed between days 2-5 (AO) and days 13-15 (EtBr). **(C)** Analysis of mitochondrial genes vs all genes analyzed in a 14-day CRISPR KO growth screen. Essential genes in K562 were defined as those with a Z<-3.5 based on sgRNA abundance between day 14 vs day 0, which includes 75% common essential genes defined by [Hart et al. 2015]. **(D)** Scatter plot shows fold change in protein abundance upon mtDNA depletion in each genotype vs that gene’s essentiality (Z score defined in C). Fraction of essential or mitochondrial genes depleted by EtBr and rescued by IF1 KO are shown in bar plots below. **(E)** Western blot for LC3B and ubiquitin phosphorylated at serine 65 on whole cell lysates of control or ATP5IF1 KO cells treated with EtBr for 12 days. Coomassie stain of total protein is shown as a loading control. **(F)** Fold change in protein abundance of all proteins in each indicated category, measured by TMT proteomics. **(G)** Relative levels of proteins in the early, essential mitochondrial Fe-S cluster biosynthesis pathway and nuclear and cytosolic Fe-S cluster-containing proteins. All measurements are made with TMT proteomics. Comparison groups for t-test: sgGFP vs sgGFP+EtBr and sgGFP+EtBr vs sgATP5IF1+EtBr. An asterisk is used to denote all significant (p value <0.05) differences. **(H)** Western blot for the 4Fe-4S cluster-containing protein POLD1 and beta tubulin for loading control. Samples are whole cell lysates of control or ATP5IF1 KO treated with EtBr for 12 days. Bar plot shows quantification of the western blots. For B, E, and H, all error bars show standard error from three biological replicates and asterisks indicate p value from a two-tailed Students t-test *<0.05, **<0.01, ***<0.001. Abbreviations: WT: wildtype, KO: knockout, EtBr: ethidium bromide, AO: antimycin and oligomycin.

Because cell proliferation and fitness directly depend on the action of essential genes, we evaluated whether a mitochondrial insult via EtBr or AO disproportionately affects essential proteins. First, we note that MitoCarta3.0^1^ is enriched for essential genes – while mitochondrial genes comprised 9.3% of all genes in our pooled CRISPR KO library, they comprised 18.8% of genes deemed as essential based on guide RNA dropout in a growth screen on day 14 (essentiality cutoff = Z score_day 14 vs day 0 < −3.5, Figure 4C)^2^. TMT proteomics of control and ATP5IF1 KO cells treated with EtBr revealed that EtBr decreased many essential proteins, all of which were rescued by ATP5IF1 loss (Figure 4D).

We hypothesized that ATP5IF1 KO, which is known to boost the PMF, rescues essential mitochondrial processes in the face of EtBr by directly rescuing mitochondrial turnover. Indeed, both autophagy marker LC3B-II and mitophagy/PINK1 activation marker phospho-S65 ubiquitin were induced by EtBr but rescued by ATP5IF1 loss (Figure 4E). Consistent with the rescue of mitochondrial turnover, ATP5IF1 KO rescued the mitochondrial matrix and inner membrane proteome in EtBr-treated cells (Figure 4F). In contrast, the ATF4 response still largely persisted in mtDNA-depleted ATP5IF1 KO cells (Figure S4B).

We next focused on the impact of mtDNA depletion on the single most conserved mitochondrial process – iron sulfur (Fe-S) cluster biosynthesis. Levels of multiple proteins in the early mitochondrial Fe-S synthesis pathway are decreased with EtBr and rescued by ATP5IF1 loss. If this dampening and rescue of the mitochondrial Fe-S synthesis machinery is biologically relevant, we would expect net Fe-S cluster insufficiency and resulting lower steady-state levels of Fe-S cluster-containing proteins^35^. Indeed, we observed a quantitative loss of all detected nuclear and cytosolic Fe-S cluster-containing proteins (except GLRX2) by EtBr and their rescue by ATP5IF1 KO (Figure 4G). Western blot validation of EtBr-induced decrease in the nuclear Fe-S cluster-containing POLD1 and its rescue by ATP5IF1 loss is shown and quantified in Figure 4H.

### mtCN setpoint is under opposing regulation by cell proliferation and mtDNA replication, both of which require the PMF

Building on the genetic evidence from ATP5IF1 KO, we directly modulated the PMF pharmacologically and tested its impact on cell proliferation and mtCN simultaneously. We treated K562 cells with a small molecule uncoupler BAM15 to decrease the PMF. BAM15 treatment slowed both – cell proliferation rate as well as mtDNA replication rate – in a dose-dependent manner, but the steady state mtDNA copy number setpoint was unchanged even at the highest tested dose of BAM15 (Figure 5A).

**Figure 5.**
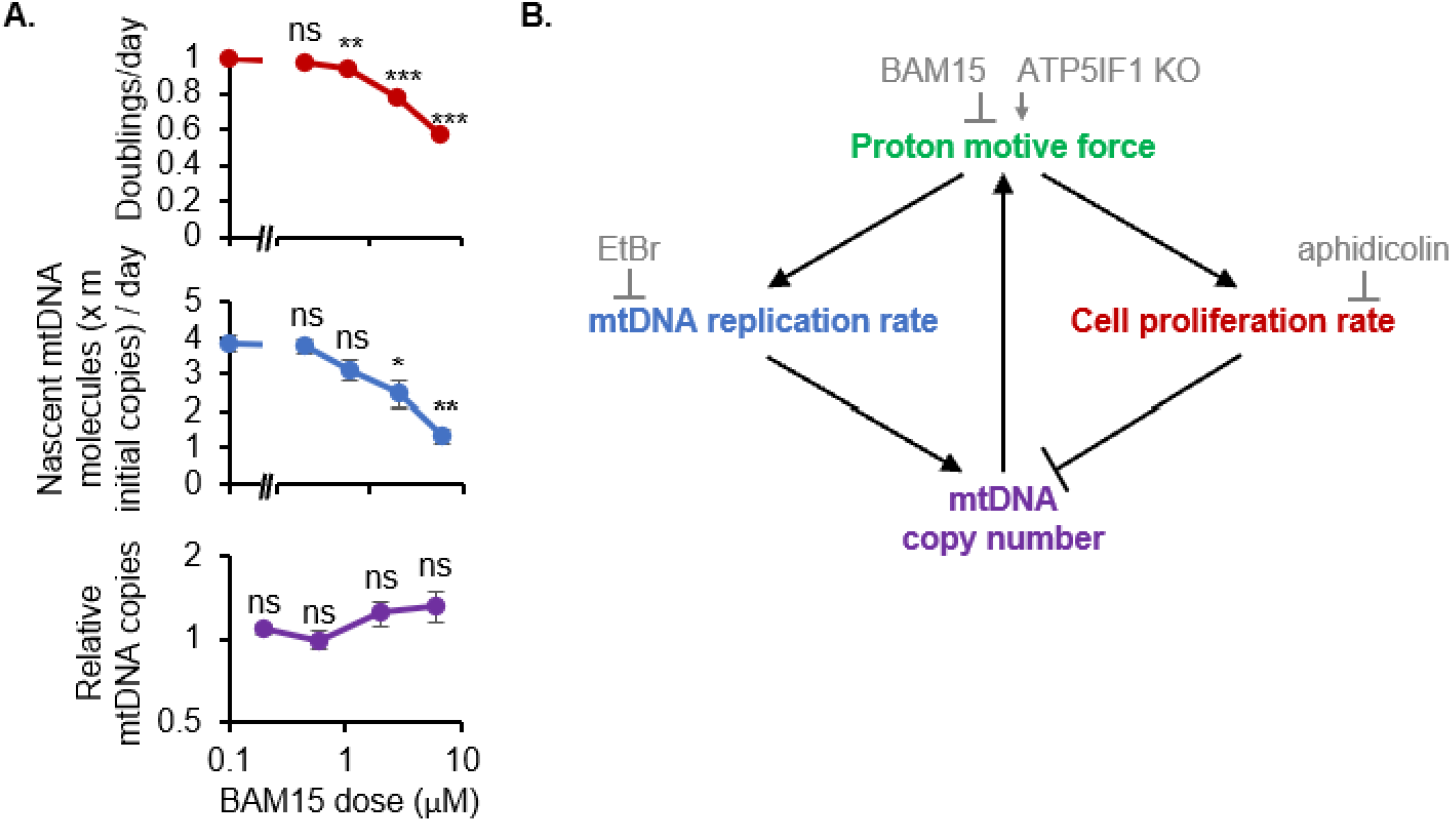
Model for the regulation of mtDNA copy number by the proton motive force in dividing cells. **(A)** Effects of uncoupler BAM15 on cell proliferation, the calculated rate of mtDNA replication, and steady state mtDNA copy number measured by qPCR. Error bars represent standard error of three replicates and asterisks indicate p value from a two-tailed Students t-test *<0.05, **<0.01, ***<0.001. **(B)** Model showing that the proton motive force (PMF) supports both mtDNA replication and cell proliferation, which have opposing effects on steady state mtDNA copy number. MtDNA in turn directly supports the PMF by encoding the electron transport chain. In grey are inhibitors or genetic perturbations used to probe each parameter throughout this work.

A model for mtDNA copy number regulation in dividing cells is shown in Figure 5B: The PMF boosts mtDNA replication rate and thus positively regulates mtDNA copy number. However, the PMF also caps the achievable mtDNA copy number by supporting cell proliferation, which dilutes mtDNA copies by half with each cell doubling. Cell proliferation is thus a built-in brake on mtDNA copy number. The PMF supports cell proliferation at least in part by preventing mitophagy and allowing mitochondrial import/stability of proteins that execute essential cellular functions such as Fe-S cluster biosynthesis. This model emerges directly from perturbative experiments presented in this work which modulate each node: EtBr (to inhibit mtDNA replication rate), aphidicolin (to inhibit cell proliferation rate), and ATP5IF1 KO and BAM15 (to boost and inhibit the PMF respectively). Altogether, the PMF is necessary for both mtDNA replication and cell proliferation – the opposing effects of which determine the mtDNA copy number setpoint of dividing cells.

## Discussion

In this work, we leverage the mtDNA depletion-recovery system to investigate how K562 cells sense and regulate their mtCN. Our proteomic, ultrastructure, and bioenergetic studies reveal that cells support mtCN recovery despite pervasive mitochondrial damage and a largely abolished ETC. In this regime, residual ETC activity and CV reversal defend the PMF, which is required for mtCN recovery. The ETC can also be completely dispensable for mtCN recovery with uridine salvage and unhindered complex V reversal enabled by ATP5IF1 KO, allowing cells to escape a “circular dependence” of mtDNA on the mtDNA-encoded ETC. We discover that cell proliferation is not only impaired by mtDNA depletion but, when it resumes upon mtCN recovery, proliferation is a major dilutive force that restricts the achievable per-cell mtCN. A delay in resuming proliferation often leads to a modest mtCN overshoot and chemically inhibiting proliferation altogether further boosts mtCN. We probe how proliferation is coupled to mtCN by examining the sole, conserved suppressor (ATP5IF1) whose KO is long known to rescue the proliferation of mtDNA-depleted cells via the PMF. We show that mtDNA depletion collapses the mitochondrial proteome, and with it, several essential processes as evidenced by Fe-S cluster starvation – all of which is rescued by ATP5IF1 KO. Thus, proliferation is coupled to mtCN via essential, mitochondria-dependent processes, supported by the PMF. Hence, PMF supports both mtDNA replication and cell proliferation, whose opposing effects determine a cell’s mtCN setpoint.

Our temporal multi-omic and bioenergetic analyses of mtDNA depletion-recovery pinpoint that mtDNA recovery occurs in the face of pervasive mitochondrial damage and ETC deficiency. We further show that the mtDNA-encoded ETC can be completely dispensable for mtDNA replication as long as cells can defend their PMF by unhindered complex V reversal and salvage pyrimidines. This is consistent with three prior observations: (1) patients harboring ETC defects do not universally exhibit mtDNA depletion^36^, (2) mtDNA deletion/depletion syndromes (MDDS) are a distinct class of mitochondrial disorders that typically do not arise from mutations in the ETC^37^, and (3) the ETC did not score in genetic screens for regulators of mtCN^38^. On the other hand, the PMF and pyrimidines are previously linked to mtDNA maintenance –mitochondrial uncoupler was shown to inhibit nascent mtDNA synthesis in drosophila oogenesis^39^, and mutations leading to nucleotide pool imbalance are linked to MDDS^29^.

To understand how the ETC can be dispensable for human mtDNA maintenance, we can consider what is strictly required for mtDNA replication: the mtDNA replication machinery, purine synthesis/salvage, pyrimidine synthesis/salvage, and the PMF. Of these, the first two are entirely “nuclear-encoded,” while the latter two can depend either on the combination of mtDNA-encoded and nuclear genes (CIII-dependent DHODH and CI-III-IV-driven proton pumping, respectively) or exclusively on nuclear genes (pyrimidine salvage and CV reversal respectively). Our results indicate that mtDNA recovery can be sustained exclusively by a “nuclear solution” for pyrimidine salvage and PMF. However, this bioenergetic flexibility is not apparent in other eukaryotes: for example, in Amoebozoa, the ETC and the alpha subunit of CV (atp1) which participates in CV reversal are both mtDNA-encoded^40^, which predicts that these organisms may not tolerate and recover from mtDNA loss as it would impair both the known modes of PMF generation. In contrast, in trypanosoma, where like metazoa and fungi, the F1 is exclusively nuclear-encoded, and it is known these organisms rely on CV reversal to survive stages of their life cycle that are mtDNA-depleted^41^. It is tempting to speculate that re-localization of the genes required for the F1-ATPase may have evolved to create a flexibility that allows overcoming the circular dependence between mtDNA and the mtDNA-encoded ETC.

We home in on a striking feature of the mtDNA depletion-recovery system – the fact that mtCN returns to its initial baseline after a mild overshoot – and demonstrate that the reciprocal regulation of mtCN & cell proliferation is directly responsible for this phenomenon. First, cell proliferation slows in response to mtDNA depletion & remains stalled during recovery, which prevents further mtCN dilution and enables mtCN recovery. Then, only after full mtCN recovery does the ETC re-engage and stress responses such as the ISR & PINK1 activation resolve, concordant with which proliferation resumes and offsets the mtDNA replication rate – this re-establishes the original mtCN setpoint. We speculate that the slight delay in cells resuming proliferation leads to the observed overshoot of mtCN and mtDNA-encoded proteins, suggesting that proliferation serves as a brake on mtCN. Indeed, we observe that inhibiting proliferation altogether increases mtCN in a depletion-recovery experiment as well as the basal mtCN of all four human cell lines tested. This is in line with previous reports that mtDNA replication occurs throughout the cell cycle^42^ and is unaffected by cell cycle inhibitors. A recent study in budding yeast showed that G1 arrest increases mtCN concordant with cell volume^43^. Genetic perturbations reported to increase mtCN typically over-express proteins involved in mtDNA replication, including POLG^44^, TFAM^44,45^,46, Twinkle^47^, TFB2M^48^, and SSBP1^49^ suggesting that they are limiting for mtCN. Thus, likely a combination of such “limiting factors” and dilution by cell proliferation restricts the per cell mtCN.

To delineate how proliferation is coupled to mtCN, we characterize the proliferation rescue of mtDNA-depleted cells by the sole and conserved suppressor, ATP5IF1 KO. We explain this proliferation rescue via the lens of essentiality: MitoCarta is enriched for essential proteins and ATP5IF1 KO rescues the mitochondrial proteome, including the Fe-S cluster biosynthesis machinery and in turn Fe-S cluster starvation. Fe-S clusters are indispensable in numerous cellular processes including nuclear genome replication^50^. In line with this, mitochondrial dysfunction is reported to cause nuclear genome instability in yeast due to Fe-S cluster starvation^51^. We therefore propose that mtDNA, which is normally responsible for generating the PMF via the ETC, directly licenses cell proliferation through the provision of mitochondria-dependent essential factors, including Fe-S clusters.

## Methods

### Cell culture

All cell lines used in this study were cultured in DMEM (Gibco 11995-065) with 10% FBS (Sigma Aldrich F2442), and 100 U/mL penicillin/ streptomycin under 5% CO_2_ at 37°C. The ATP5IF1 KO polyclonal K562 cell line was generated using two guide RNAs against ATP5IF1 together: CGGACGTGGCTTGGCGTGTG and GTGAGGACCATGCAAGCCCG; the control cell line was generated using a guide RNA against EGFP (GGGCGAGGAGCTGTTCACCG.

### mtDNA depletion-recovery assay and mtDNA quantification

Throughout the timecourse, K562 cells were cultured in DMEM (Gibco 11995-065) with 10% dialyzed FBS (Gibco 26400044), 100 U/mL penicillin/streptomycin, and 50 μg/mL supplemental uridine. MtDNA was depleted by adding 100 ng/mL EtBr for the first five days and mtCN recovery was enabled by transferring the mtDNA-depleted cells to EtBr-free media for 15-20 days. Cells were passaged every 2-3 days, diluting to a confluence of 1-2×10^5^ cells/mL and harvested at various timepoints during the depletion-recovery timecourse for analyses of mtCN, whole cell proteome, ultrastructure, bioenergetics, and proliferation.

To quantify relative mtCN, total genomic DNA was extracted using Qiagen DNeasy kit following the manufacturer’s protocol and 30 ng DNA was used for each qPCR reaction. Two mitochondrial probes were each normalized to a nuclear probe to calculate mtDNA copy number relative to nuclear genome copies. All mtCNs were plotted as fold change relative to day 0 in the depletion-recovery timecourse, and relative to no aphidicolin in Fig 3C, using the ΔΔCt method. Average of both mitochondrial probes across three biological replicates is reported. Primers used were:

mt-tRNA-Leu1 forward: CACCCAAGAACAGGGTTTGT

mt-tRNA-Leu1 reverse: TGGCCATGGGTATGTTGTTA

mt-ND6 forward: CCACACCGCTAACAATCAATAC

mt-ND6 reverse: GTTTCTGTTGAGTGTGGGTTTAG

nuclear B2M forward: GCTGGGTAGCTCTAAACAATGTATTCA

nuclear B2M reverse: CCATGTACTAACAAATGTCTAAAATGGT

### TMT proteomics and phosphoproteomics

Quantitative proteomics was performed at the Thermo Fisher Scientific Center for Multiplexed Proteomics (Harvard).

Sample preparation for mass spectrometry: Cell pellets were lysed in 8M urea, 200 mM EPPS pH 8.5, protease and phosphatase inhibitors. Proteins were quantified using a Pierce micro-BCA assay. Samples were then reduced with TCEP, alkylated with iodoacetamide, and precipitated by methanol/chloroform. Precipitated proteins were resuspended in EPPS, digested with LysC (1:50 protease:protein) and trypsin (1:100). Peptides were quantified and completely labeled with TMT11 reagents (Thermo-Fisher).

For phosphoproteomics, pSTY peptides were enriched from the combined sample using the Pierce High-Select Fe-NTA phosphopeptide enrichment kit. The unbound peptides were recovered, cleaned, dried by speedvac, and fractionated by HPLC. pSTY peptides were eluted, desalted, dried by speedvac, cleaned on a stage tip, and eluted into an MS glass sample vial. The phos-enriched sample was analyzed twice by LC-MS3 on an Orbitrap Lumos Mass Spectrometer equipped with a FAIMS ion source using a 180-minute method. Fragmentation was achieved in one run by MSA and the other run by HCD.

Basic pH reverse phase separation: TMT labeled peptides were solubilized in 5% ACN/50 mM ammonium bicarbonate, pH 8.0 and separated by an Agilent 300 Extend C18 column (3.5μm particles, 4.6 mm ID and 250 mm in length). An Agilent 1260 binary pump coupled with a photodiode array (PDA) detector (Thermo Scientific) was used to separate the peptides. A 60 min linear gradient from 10% to 90% acetonitrile in 50 mM ammonium bicarbonate pH 8.0 separated the peptide mixtures into a total of 96 fractions (fraction size – 0.37 min, 300 uL). These fractions were consolidated into 24 total samples in a checkerboard fashion, dried by speedvac, cleaned on a C18 packed stage tip, and re-dissolved in 5% Acetonitrile/5% formic acid/ and a third of the sample was analyzed by LC-MS3.

Liquid chromatography separation and tandem mass spectrometry: 12 fractions from the total proteome were analyzed on an Orbitrap Lumos mass spectrometer equipped with a FAIMS ion source using a 180 minute MS3 method. Peptides were detected (MS1) and quantified (MS3) in the Orbitrap, and sequenced (MS2) in the ion trap.

Data analysis: MS2 spectra were searched using the SEQUEST algorithm against a Uniprot composite database derived from the human proteome containing its reversed complement and known contaminants. Peptide spectral matches were filtered to a 1% false discovery rate (FDR) using the target-decoy strategy combined with linear discriminant analysis. The proteins from the 12 runs were filtered to a <1% FDR. Proteins were quantified only from peptides with a summed SN threshold of >=100. For phosphosites, MS2 spectra were searched using the SEQUEST algorithm against a Uniprot composite database described previously with phosphorylation on Ser, Thr, or Tyr as differential modifications. Phosphopeptide FDR was filtered to 1% by LDA analysis of modified peptides only. Proteins were filtered to 1% FDR. Phosphosite localization was scored by ModScore and proteins were quantified using peptides with a summed signal-to-noise threshold of >=100. To control for different total protein loading within a TMT experiment, the summed protein quantities of each channel were adjusted to be equal within the experiment. Proteins quantified by >1 peptide were analyzed. Fold change in each protein relative to day 0 was calculated across the depletion-recovery timecourse.

### Transmission electron microscopy

Non-adherent K562 CML cells, grown in 10 cm dishes, were fixed with 3% Glutaraldehyde for 2-3 hours at room temperature, then allowed to infiltrate in fixative overnight at 4 °C. Cell suspensions were pelleted (4500 rpm 20 min at 4°C), rinsed several times in 0.1M sodium cacodylate buffer, post-fixed 1 hour in 1% OsO 4 /0.1M cacodylate buffer, then rinsed again several times in buffer, re-centrifuging between steps as needed to keep pellet aggregate together. Pellets were embedded in 2% agarose, dehydrated through a graded series of ethanols to 100%, briefly dehydrated in 100% propylene oxide and allowed to infiltrate overnight on a gentle rotator in a 1:1 mix of propylene oxide and Eponate resin (Ted Pella, Redding, CA). The following day, samples were transferred into freshly prepared 100% Eponate resin and allowed to infiltrate several hours at room temperature. Agarose blocks were embedded in flat molds in 100% fresh Eponate resin and placed into a 60°C oven for polymerization (24-48 hours). Thin (70 nm) sections were cut using a Leica EM UC7 ultramicrotome, collected onto formvar-coated copper slot grids (Electron Microscopy Sciences, Hatfield, PA), stained with 2% uranyl acetate and Reynold’s lead citrate and examined in a JEOL JEM 1011 transmission electron microscope at 80 kV. Images were collected using an AMT digital imaging system with proprietary image capture software (Advanced Microscopy Techniques, Danvers, MA).

### Seahorse MitoStress Test and ATP rate assay

Multiple mtDNA depletion-recovery timecourses were staggered such that the desired timepoints along the timecourse were available for Seahorse and ATP rate assay on a single day. Both assays were performed as per manufacturer’s instructions using 1×10^5^ cells/well in XF DMEM pH 7.4 (Agilent 103574-100), 10 mM glucose (Agilent 103577-100), 1 mM pyruvate (Agilent 103578-100) and 4 mM glutamine (Gibco 25030-081). Final drug concentrations after injection into the well were 1.5 μM oligomycin, 0.8 μM CCCP, and 1.5 μM antimycin and piericidin. Data were analyzed using Seahorse’s report generator Excel macro (Agilent).

### Membrane potential measurement

Naïve or mtDNA-depleted K562 cells were incubated with 10 nM TMRM in the presence of either DMSO, 100 nM antimycin, 30 nM oligomycin, or 100 nM antimycin and 30 nM oligomycin for 30 mins at 37°C under 5% CO_2_. Cells were then quickly washed with PBS and 3×10^5^ cells were plated per well in 96-well plates to measure emission at 573 nm using a platereader (2104 Multilable Reader Envision, Perkin Elmer).

### Proliferation assay and mtDNA replication rate calculation

Viable cells were counted using Vi-CELL BLU (Agilent). Proliferation rate was calculated as Log_2_(final/initial viable cell count) divided by the number of days elapsed between timepoints. Steady state mtCN was determined using qPCR as described above and reported relative to day 0. To calculate the mtDNA replication rate, the total number of new mtDNA molecules that had to be made between two timepoints was calculated by accounting for both: (A) fold increase/decrease in steady state mtCN between these timepoints (final/initial mtDNA copies) and (B) fold dilution of mtCN by cell doubling that had to be buffered (2^number of doublings that occurred between the time interval). Relative mtDNA molecules made between timepoints = mtCN relative to day 0 at the start of the time interval x A x B.

### Cell cycle analysis

Multiple mtDNA depletion-recovery timecourses were staggered such that the desired timepoints along the timecourse were all available for cell cycle analysis on a single day. Cells were pelleted at 650 g for 2 min, washed once with PBS, and fixed in 4% paraformaldehyde for 20 mins at room temperature. Cells were then washed once with PBS, permeabilized with 0.1%Tx-100 for 10 mins, and washed again with PBS. Cells were treated with 1U DNase-free RNase for 30 mins at 37°C and then stained with 50 µg/ml propidium iodide, washed, and analyzed on a SORP 5-Laser BD LSRFortessa. Cell cycle phases were quantified and visualized using the Watson Pragmatic algorithm in FlowJo.

### Western blot

Equal number of cells from each condition (1-2 ×10^6^) were harvested by pelleting at 650 g for 2 min, washed once with PBS, and lysed directly in 150 μL LDS sample buffer (Nupage NP0007) diluted to 2x with water. Ten percent of the sample was run on Novex™ 4-20% Tris-Glycine Mini Gels (Thermo Fisher Scientific XP04200BOX) and transferred to a 0.2 μm PVDF membrane (BioRad 1704156). The gel was subsequently stained with Coomassie to stain total proteins and membranes were blocked for 10 minutes in 5% non-fat dry milk prepared in TBST. Membranes were then incubated with primary antibody diluted 1:1000 in 5% milk overnight at 4 °C. The primary antibodies used in this work are: LC3B (Abcam ab192890), phospho-S65 ubiquitin (Cell Signaling 62802S), POLD1 (Abcam ab186407). Membranes were then washed in TBST 4 times for 5 minutes each at room temperature. Next, they were incubated in HRP-linked α-rabbit (Cell Signaling 7074P2) or α-mouse (Cytiva NA931V) secondary antibodies diluted 1:5000 in 5% non-fat dry milk, for 1 hour at room temperature. Membranes were washed again in TBST as before, incubated with 2 mL of Lightning® Plus ECL (Revvity NEL103E001EA), and exposed to Amhersham Hyperfilm™ (GE Healthcare). Band intensities were quantified using GelQuant (BiochemLabSolutions) and normalized to total protein.

## Data availability

All proteomics data will be deposited to PRIDE at the time of publication.

## Code availability

There is no code associated with this study.

## Acknowledgements

We thank all members of the Mootha laboratory for helpful discussions and feedback. We thank Steve Gygi, Rachel Beth Rodrigues, and the TCMP facility at Harvard Medical School for TMT proteomics, the MGH Pathology Flow Cytometry Core for access to Flow Cytometry. For electron microscopy, we thank Diane Capen at the Microscopy Core of the Center for Systems Biology/Program in Membrane Biology, which is partially supported by an Inflammatory Bowel Disease Grant DK043351 and a Boston Area Diabetes and Endocrinology Research Center (BADERC) Award DK057521. We thank Tsz-Leung To for generating the ATP5IF1 KO cell line. This work was supported by NIH grants K00CA212468 and K99GM145848 (to S.P.R.). V.K.M. is an investigator of the Howard Hughes Medical Institute.

## Author contributions

S.P.R. designed and performed experiments and analyzed the data. S.P.R. and V.K.M. conceptualized the study and wrote the manuscript.

## Competing interests

V.K.M is a paid advisor to 5AM ventures and Falcon Bio.

## Figures

**Figure S1:**
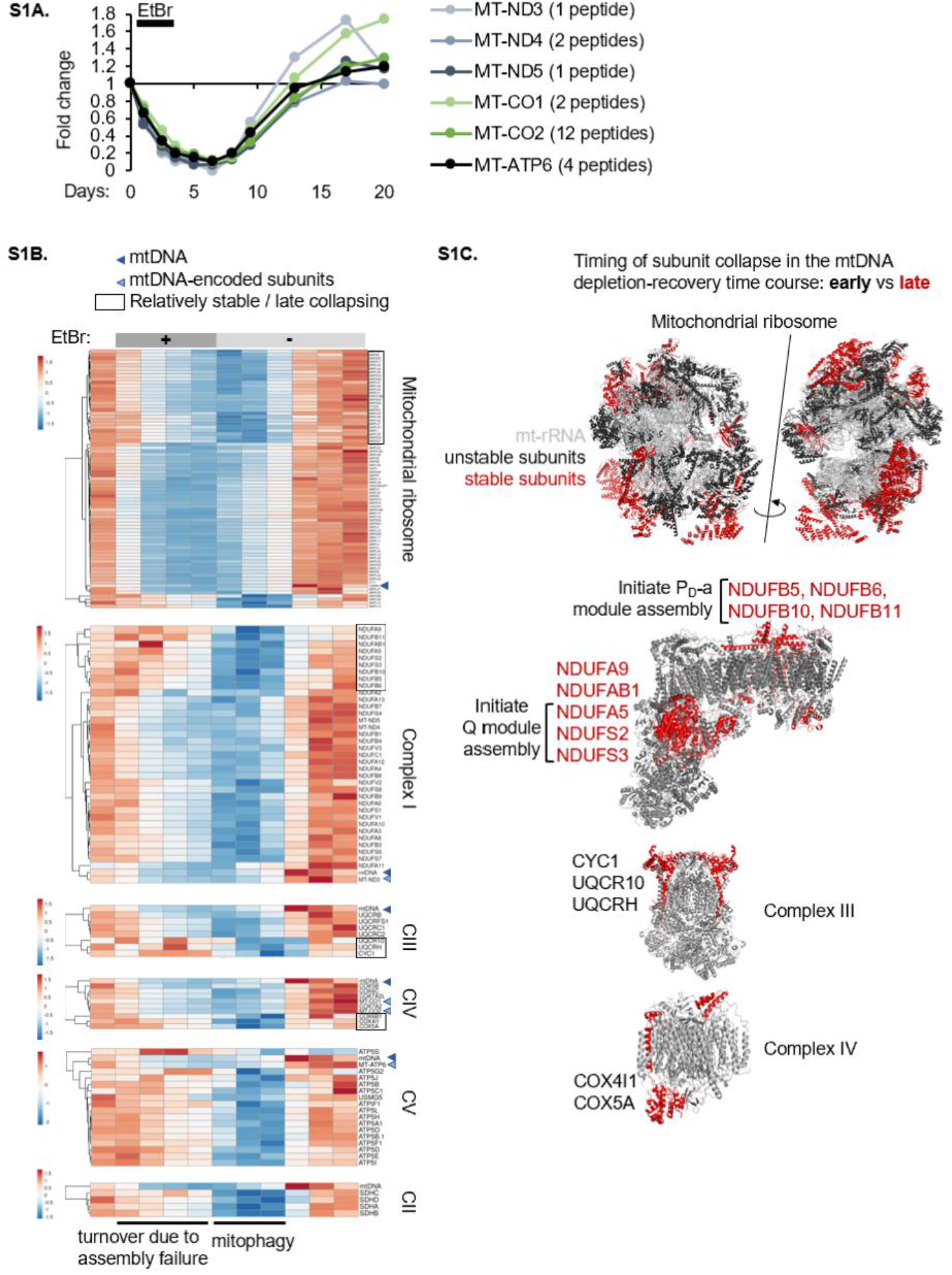
Temporal analysis of subunit loss upon mtDNA depletion in the context of the 3D structure of each complex. **(A)** Levels of mtDNA-encoded proteins detected by TMT proteomics during mtDNA depletion-recovery. **(B)** Heat map of all proteins detected in each indicated complex. mtDNA levels and mtDNA-encoded proteins are indicated by arrows. Black box indicates relatively stable subunits, which collapse later, as revealed by hierarchical clustering. **(C)** Subunits of all dual genome-encoded complexes color-coded on the 3D structure by timing of collapse in the mtDNA depletion-recovery system (early vs late collapsing).

**Figure S2.**
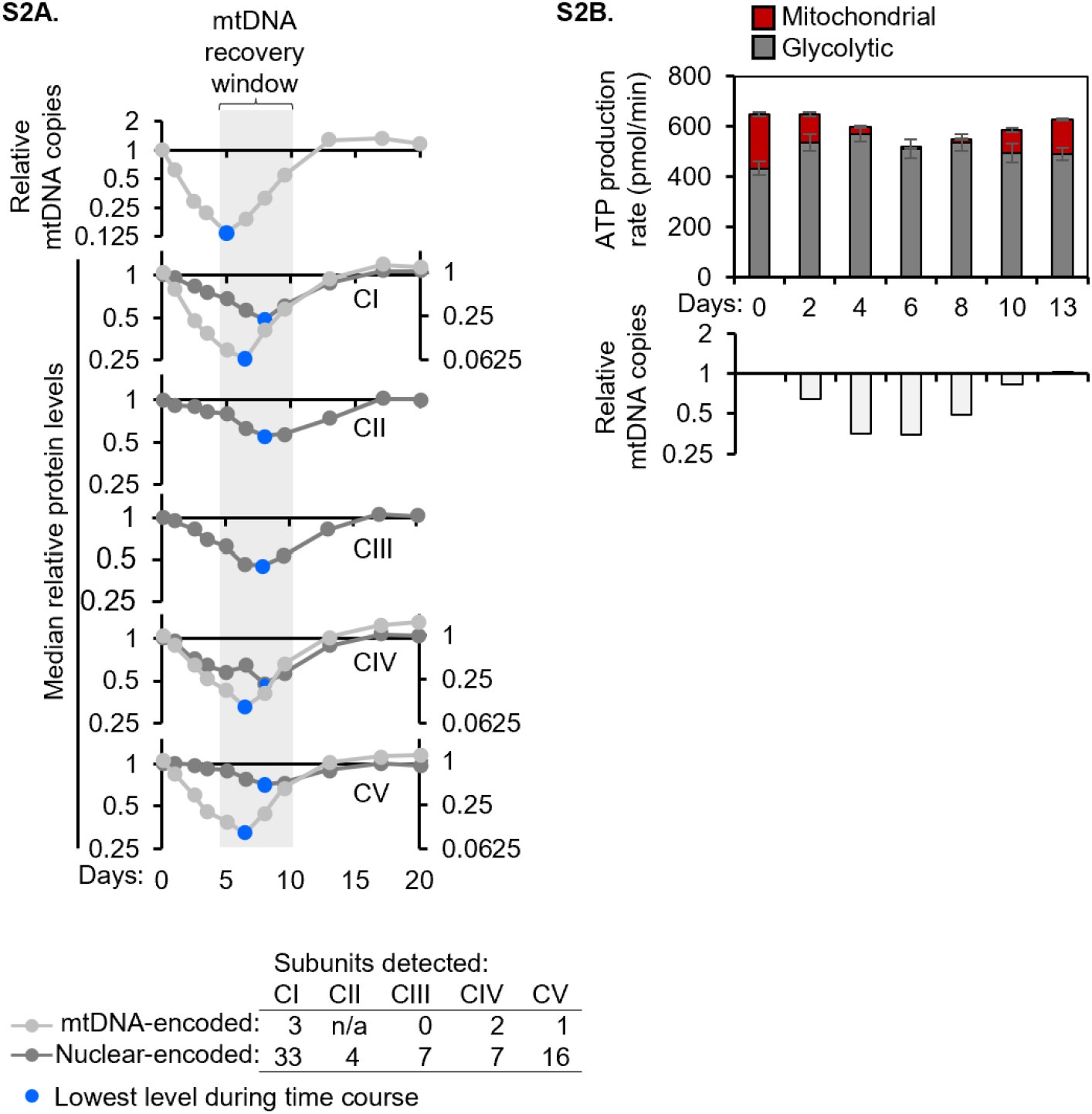
OXPHOS remains diminished during mtDNA recovery. **(A)** Median levels of all OXPHOS complex subunits detected by TMT proteomics. MtDNA- and nuclear-encoded subunits are shown separately. A blue dot is used to indicate the lowest level of each entity during the mtDNA depletion-recovery timecourse. (**B)** Stacked bar plots of mitochondrial and glycolytic ATP production rates measured in at least three replicates during a representative mtDNA depletion-recovery timecourse and mtDNA copy number measured by qPCR at the corresponding timepoints.

**Figure S3:**
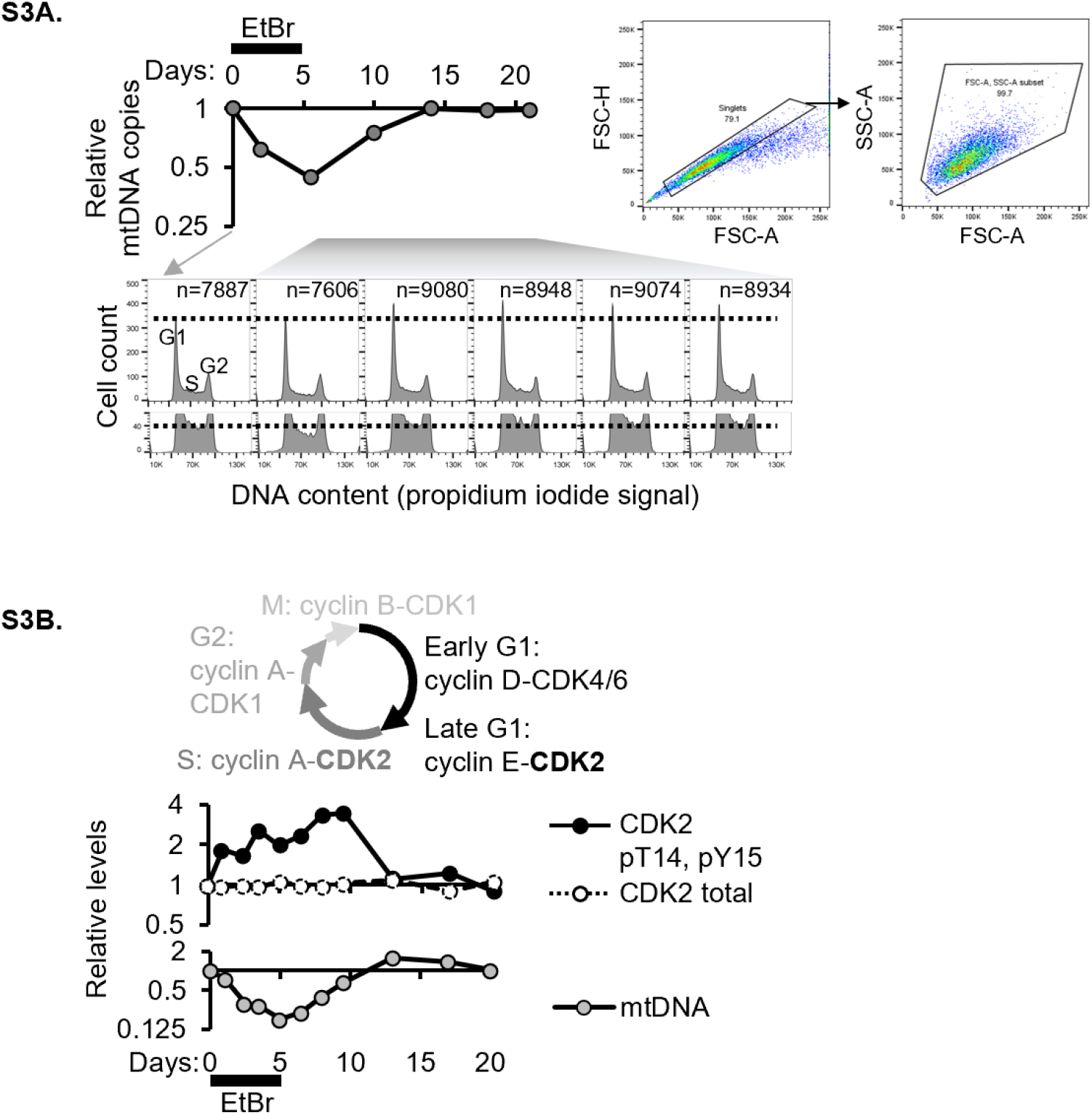
Cell cycle analysis during mtDNA depletion and recovery. **(A)** Cell cycle analysis by flow cytometry of asynchronous cultures from day 0 and all timepoints during mtDNA copy number recovery. Bottom panel is zoomed to show S phase, horizontal lines are drawn to aide visualization of the slight up-tick in G1 and slight decrease in S phase on day 5. Gates used to select single cells and specify side and forward scatter areas are shown. **(B)** Schematic of the cell cycle. Quantification of mtDNA copy number (qPCR), CDK2 protein, and CDK2 phospho-peptide capturing the inhibitory phosphosites pT14, pY15 (TMT proteomics).

**Figure S4:**
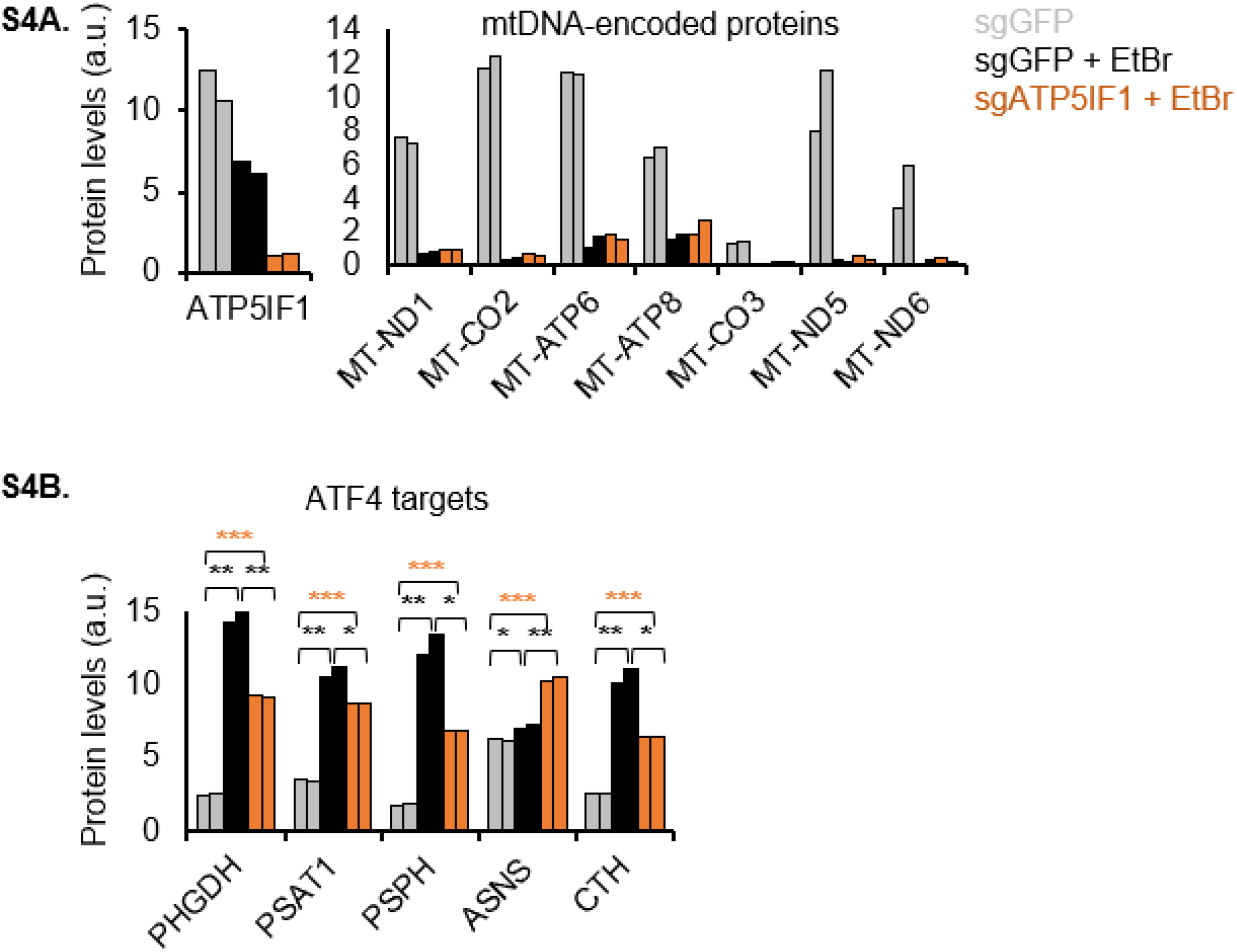
ATF4 response persists in mtDNA-depleted ATP5IF1 KO cells. Protein levels of **(A)** ATP5IF1, mtDNA-encoded proteins, and **(B)** ATF4 targets in control vs ATP5IF1 KO treated with EtBr for 14 days. Asterisks denote p value from a two-tailed Students’ t-test *<0.05, **<0.01, ***<0.001.

